# Identifying highly-penetrant disease causal mutations using next generation sequencing: Guide to whole process

**DOI:** 10.1101/011130

**Authors:** Mesut Erzurumluoglu

**Affiliations:** Bristol Genetic Epidemiology Laboratories (BGEL), School of Social and Community Medicine, University of Bristol, Oakfield House, Oakfield Grove, Bristol BS8 2BN, United Kingdom.

## Abstract

Recent technological advances have created challenges for geneticists and a need to adapt to a wide range of new bioinformatics tools and an expanding wealth of publicly available data (e.g. mutation databases, software). This wide range of methods and a diversity of file formats used in sequence analysis is a significant issue, with a considerable amount of time spent before anyone can even attempt to analyse the genetic basis of human disorders. Another point to consider is although many possess ‘just enough’ knowledge to analyse their data, they do not make full use of the tools and databases that are available and also do not know how their data was created. The primary aim of this review is to document some of the key approaches and provide an analysis schema to make the analysis process more efficient and reliable in the context of discovering highly penetrant causal mutations/genes. This review will also compare the methods used to identify highly penetrant variants when data is obtained from consanguineous individuals as opposed to non-consanguineous; and when Mendelian disorders are analysed as opposed to common-complex disorders.

## INTRODUCTION

Next generation sequencing (NGS) and other high throughput technologies have brought new challenges concomitantly. The colossal amount of information that is produced has led researchers to look for ways of reducing the time and effort it takes to analyse the resulting data whilst also keeping up with the storage needs of the resulting files – which are in the magnitude of gigabytes each. The recently emerged variant call format (VCF) has somewhat provided a way out of this complex issue [1]. Using a reference sequence and comparing it with the query sequence, only the differences between the two are encoded into a VCF file. Not only are VCF files substantially smaller in size (>300x in relation to BAM files which store all raw read alignments), they also make the data relatively easy to analyse since there are many bioinformatics tools (e.g. annotation, mutation effect prediction) which accept the VCF format as standard input. The Genome Analysis Toolkit (GATK) made available by the Broad Institute also provides useful suggestions to bring a universal standard for the annotation and filtering of VCF files [2]. The abovementioned reasons have made VCF the established format for the sharing of genetic variation produced from large sequencing projects (e.g. 1000 Genomes Project, NHLBI Exome Project - also known as EVS). However the VCF does have some disadvantages. The files can be information dense, initially difficult to understand and parse. Comprehensive information about the VCF and its companion software VFCtools [1] are available online (vcftools.sourceforge.net).

Because of the substantial decrease in the price of DNA sequencing and genotyping [3], there has been a sharp increase in the number of genetic association studies being carried out, especially in the form of genome-wide association studies (GWAS, statistics available at www.genome.gov/gwastudies/). As whole genome sequencing (WGS) is prohibitively expensive for large genetic association studies [4-6], whole exome sequencing (WES) has emerged as the attractive alternative – where only the protein coding region of the genome (i.e. exome) is targeted and sequenced [7]. This decision to carry out WES over WGS is not solely influenced by the cost which currently stands at one-third in comparison [8], but also by the fact that most of the known Mendelian disorders (∼85%) are caused by mutations in the exome [9] and reliably interpreting variation outside of the exome is still challenging as there is little consensus (even with ENCODE data [10] and non-coding variant effect prediction tools such as CADD [11] and GWAVA [12]). For complex diseases, WES can provide more evidence for causality compared to GWAS, assuming that the causal variants are exonic. This is because the latter uses linkage disequilibrium (LD) patterns between common markers [13] whereas WES directly associates the variant itself with the phenotypes/disorder. Therefore using GWAS, especially in gene-dense regions, one cannot usually make conclusive judgements about which gene(s) is causal without further sequencing or functional analysis. WES has been successfully used in identifying and/or verifying over 300 causal variants for Mendelian disorders (statistics from omim.org/) [14,15]. WES currently stands at approx. $1000 for 50x read depth (variable prices, less for larger studies). However since there is a great deal of variation in the human genome [16], finding the causal variant(s), especially ones with low penetrance, is not going to be trivial. This problem can be exacerbated by the nature of the disorder(s) analysed. It is relatively easier to map variants causing rare monogenic diseases, as there is most likely to be a single variant present in the cases that is not in the controls; but in contrast, common complex (polygenic) disorders are much harder to dissect when searching for causal variants.

In this paper, our aims are to (i) provide a guide for genetic association studies dealing with sequencing data to identify highly penetrant variants (ii) compare the different approaches taken when data is obtained from unrelated or consanguineous individuals, and (iii) make suggestions about how to rank single nucleotide variation (SNV) and/or insertion/deletions (indels) following the standard filtering/ranking steps if there are several candidate variants. To aid the process of analysing sequencing data obtained from consanguineous individuals, we have also made available an autozygosity mapping algorithm (AutoZplotter) which takes VCF files as input and enables manual identification of regions that have longer stretches of homozygosity than would be expected by chance.

## STAGE 1 - QUALITY CONTROL & VARIANT CALLING

Before any genetic analysis, it is important to understand how the raw data were produced and processed to make better judgements about the reliability of the data received. Thorough quality control steps are required to ensure the reliability of the dataset. Lack of adequate prior quality control will inevitably lead to loss of statistical power; and increase false positive and false negative findings. Fully comprehending each step during the creation of the dataset will have implications on the interpretation stage, where genotyping errors (also known as ‘phantom’ mutations [17]) may turn out to be statistically associated (e.g. batch effects between case and control batches) or the causal variant may not be identified due to poorly applied quality control (QC) and/or filtering methods. The most fitting example for this comes from a recent Primary ciliary dyskinesia (PCD) study [18], where the causal variant was only detected after the authors manually noticed an absence of reads in the relevant region of the genome (personal communication with authors). The subsequent variant was not only missing in the VCF files, but also in the initial BAM files - requiring remapping of reads. Another point of consideration from this finding would be that the authors knew where to look because the *RSPH9* gene (and the p.Lys268del mutation) was one of their *a priori* candidates [19]. This is also an example demonstrating the importance of deep prior knowledge and screening for known variants as it is impossible for one to manually check the whole exome (or the genome) for sequencing and mapping errors.

### Targeted sequencing

As far as WES projects are concerned, questions about coverage arise right from the start (Figure 1). Since knowledge concerning exons in our own genome is far from complete, there are differing definitions about the human exome coordinates. Therefore, the targeted regions by the commercially available Agilent SureSelect [20] and the Nimblegen SeqCap EZ [21] exome capture kits are not entirely overlapping [22]. Thus it is possible that the missing regions of the exome due to the chosen probe kit may turn out to have the functional region in relation to the disorder analysed. One must also bear in mind that the kits available for targeting the exome are not fully efficient due to a certain quantity of poorly synthesized and/or designed probes not being able to hybridize to the target DNA. Next step is target enrichment where high coverage is vital as NGS machines produce more erroneous base calls compared to other techniques [23]; therefore, especially for rare variant analyses, it is important to have data with high average read depth (i.e. ≥ 50x).

**Figure 1:**
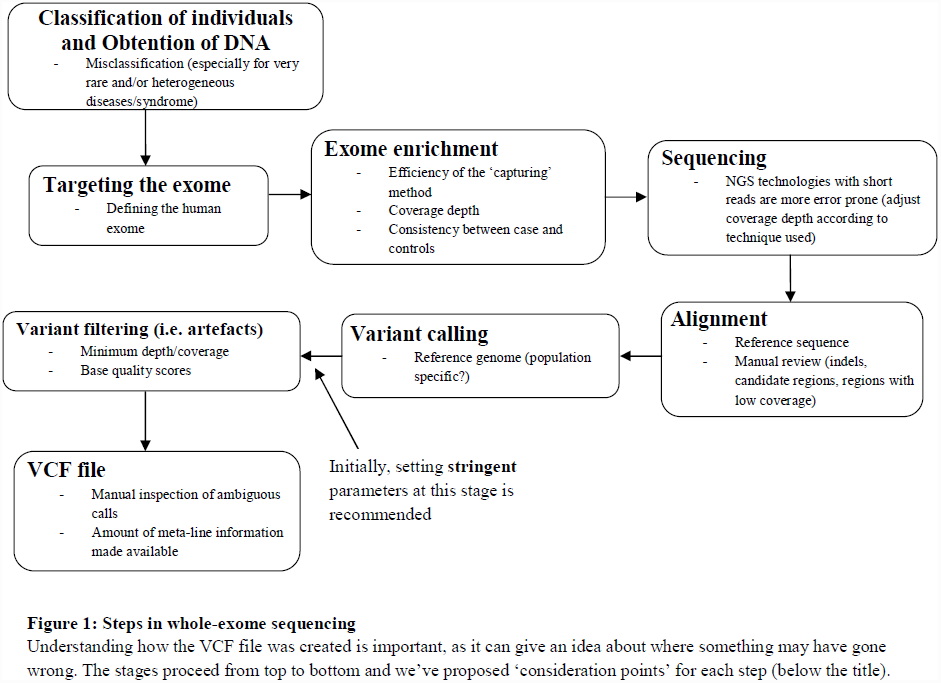
Steps in whole-exome sequencing.

### Mapping sequence reads

The raw reads produced should then be aligned to a reference genome (e.g. GRCh38 – see NCBI Genome Reference Consortium) and there are many open source and widely applied tools (Table 1). However, solely depending on automated methods and software can leave many reads spanning indels misaligned, therefore post-reviewing the data for mismapping is always a good practice, especially in the candidate regions. Attempting to remap misaligned reads with a lower stringency using software such as Pindel would be an ideal way to go about solving such a problem [24]. GATK provides a base recalibration and indel realignment algorithm for this purpose.

**Table 1:** Tools for aligning reads to a reference genome.

| **Name** | **References** | **Comment** |
| --- | --- | --- |
| BFAST | [69] | These aligners use similar algorithms to determine contiguous sequences however MAQ and BWA are widely used and have been praised for their computational efficiency and multi-platform compatibility [74]. |
| Bowtie 2 | [70] |  |
| BWA | [71] |  |
| MAQ | [72] |  |
| SOAP2 | [73] |  |

These are some of the many tools built for aligning reads produced from high throughput sequencing. Some have made speed their main purpose whereas others have paid more attention to annotating the files produced (such as mapping quality). Thus a manual review of candidate regions may prove to be crucial especially when dealing with very rare disorders.

Effective variant calling depends on accurate mapping to a dependable reference sequence. If available, using a population specific reference genome would be most ideal to filter out known neutral SNPs existing within the region of origin of the analysed subjects (e.g. East-Asian reference for subjects of Japanese origin). Inclusion of ambiguity codes (e.g. IUPAC codes) for known poly-allelic variants to create a composite reference genome can also be useful (although not essential).

### Variant calling

There are many tools available for the identification of SNVs, indels, splice-site variants and CNVs present in the query sequence(s). Each variant calling tool has advantages and disadvantages and has made compromises relating to issues such as speed of analysis, annotation and reliability of the output file (Table 2). Separating true variation from sequencing artefacts still represents a considerable challenge. When dealing with very rare disorders, the candidate regions in the output VCF (or BAM) files should be reviewed either by reviewing the QC scores in the VCF or by visualising the alignments in IGV [25]. Performing this step could highlight sequencing errors such as over-coverage (due to greater abundance of capture probes for the region or double capturing due to poorly discriminated probes hybridising to the same region) or under-coverage (due to probes not hybridising because of high variability in the region). For rare Mendelian disorders, since there is going to be a single causal variant it is important to analyse variants which are reliable. Therefore setting strict parameters for read depth (e.g. ≥ 10x), base quality score (e.g. ≥ 100) and genotype quality scores (e.g. ≥ 100) initially can eliminate wrong base and genotype calls. This can then be adjusted subsequently if no variants with a strong candidacy are found after filtering (also see Best Practices section of GATK documentation for variant analysis).

**Table 2:** Tools for identifying variation from a reference genome using NGS reads.

| Name | References | URL | Comment |
| --- | --- | --- | --- |
| GATK | [2] | http://www.broadinstitute.org/gatk/ | - Probably the most established genome analysis toolkit<br>- Includes tools such as Unified Genotyper (SNP/genotype caller), Variant filtration (for filtering SNPs) and Variant Recalibrator (for SNP quality scores)<br>- Very well documented with forums<br>- Input: SAM format<br>- Output: VCF format |
| QCALL | [75] | ftp://ftp.sanger.ac.uk/pub/rd/QCALL | - Theoretically calls 'high quality' SNPs even from low-coverage sequencing data<br>- Makes use of linkage disequilibrium information |
| PyroBayes | [76] | http://bioinformatics.bc.edu/marthlab/wiki/index.php/PyroBayes | - Theoretically makes 'confident' base calls even in shallow read coverage for reads produced by Pyrosequencing machines. |
| SAMTools | [26] | http://samtools.sourceforge.net/ | - Computes genotype likelihoods<br>- BCFtools calls SNP and genotypes<br>- Successfully used in many WGS and WES projects such as the 1000 Genomes Project [16].<br>- Offers additional features such as viewing alignments and conversion of SAM to a BAM format |
| SOAPsnp | [77] | http://soap.genomics.org.cn/soapsnp.html | - Part of the reliable SOAP family of bioinformatics tools<br>- Well documented website; and cited and used by many [78,79]. |
| Control-FREEC | [80] | http://bioinfo-out.curie.fr/projects/freec/ | - Identifies copy number variations (CNV) between case and controls from sequencing data<br>- R script available for visualising CNVs by chromosome<br>- Input format: BAM |
| Atlas2 | [81] | https://www.hgsc.bcm.edu/software/atlas-2 | - Calls SNPs and indels for WES data<br>- Requires BAM file as input<br>- Output: VCF format |

GATK, SOAPsnp and SAMTools have constantly been cited in large genetic association projects indicating their ease of use, reliability and functionality. However, this is also helped by the fact that they have additional features. There are other tools such as Beagle [63], IMPUTE2 [82] and MaCH [83] which have modules for SNP and genotype calling but are mostly used for their main purpose such as imputation and haplotype phasing.

There are many tools available for the identification of SNVs, indels, splice-site variants and CNVs present in the query sequence (see Table 2). GATK [2] is one of the most established SNP discovery and genome analysis toolkits, with extensive documentation and helpful forums. It is a structured programming framework which makes use of the programming philosophy of MapReduce to solve the data management challenge of NGS by separating data access patterns from analysis algorithms. GATK is constantly updated and cited, and also has a vibrant forum which is maintained continually.

SAMtools [26] is a variant caller which uses a Bayesian approach and has been used in many WGS and WES projects including the 1000 Genomes Project [16]. SAMtools also offers many additional features such as alignment viewing and conversion to a BAM file. A recent study has compared GATK, SAMtools and Atlas2 and found GATK to perform best in many settings [27]. However all three were highly consistent with an overlapping rate of ∼90%. SOAPsnp is another highly used SNP and genotype caller and is part of the reliable SOAP family of bioinformatics tools (http://soap.genomics.org.cn/).

### Additional checks of autozygosity

For data obtained from consanguineous families, confirming expected autozygosity (i.e. homozygous for alleles inherited from a common ancestor) would be an additional check worth carrying out. If the individual is the offspring of first cousins then the level of autozygosity would be near 6.25% (F=0.0625); and 12.5% (F=0.125) for offspring of double first cousins (or uncle-niece unions, see Supp. Figure S1 for a depiction of these). These values will be higher in endogamous populations (e.g. for offspring of first cousins: 6.25% + autozygosity brought about due to endogamy. See Supp. Fig. S3 for an example). Autozygosity could be checked by inspecting long runs of homozygosity (LRoH) for each individual by using tools such as Plink (for SNP chip data) [28], EXCLUDEAR (for SNP chip data) [29], AgilentVariantMapper (for WES data) [30] and AutoSNPa (for SNP chip data) [31] and dividing total autozygous regions by total length of autosomes in the human genome (can be obtained from http://www.ensembl.org/Homo_sapiens/Location/Genome). AutoZplotter (available to download in Supp. Materials) that we developed takes VCF files as input enabling easy and reliable visualisation and analysis of LRoH for any type of data (WGS, WES or SNP chip).

## STAGE 2 – FILTERING/RANKING OF VARIANTS

Once the quality control process is complete and VCF files are deemed analysis ready, the approach taken will depend on the type of disorder analysed. For rare Mendelian disorders, many filtering and/or ranking steps can be taken to reduce the thousands of variants to a few strong candidates. Screening previously identified genes for causal variants is a good starting point. Carrying out this simple check will allow the identification of the causal variant even from a single proband thus saving time and money. If no previously identified variant is found in the proband analysed, there are several steps which can be taken to identify novel mutations.

### Using prior information to rank/filter variants

Locus specific databases (see http://www.hgvs.org/dblist/dblist.html for a comprehensive list) and ‘whole-genome’ mutation databases such as HGMD [32], ClinVar [33], LOVD (www.lovd.nl/) and OMIM [34] are very informative resources for this task. Finding no previously identified variants indicates a novel variant in the proband analysed. For rare Mendelian disorders, the look for the variant can begin by removal of known neutral and/or common variants (≥ 0.1%) as this would provide a smaller subset of potentially causal variants. This is a pragmatic choice as Mendelian disease causal variants are likely to be very rare in the population or unique to the proband. If the latter is true, the variant will be absent from public databases. For this process to be thorough, an automated annotation tool such as Ensembl VEP can be used. VEP enables incorporation of MAF (or GMAF, global MAF) from the EVS and the 1000 Genomes Project (see Supp. Material and Methods for details).

### Using effect prediction algorithms to rank/filter variants

Ranking this subset of variants based on consequence (e.g. stop gains would rank higher than missense) and scores derived from mutation prediction tools (e.g. ‘probably damaging’ variants would rank higher than ‘possibly damaging’ according to Polyphen-2 prediction) would enable assessment of the predicted impact of all rare mutations. It is important to understand what is assumed at each filtering/ranking stage; and comments are included about each assumption and their caveats in Figure 2.

**Figure 2:**
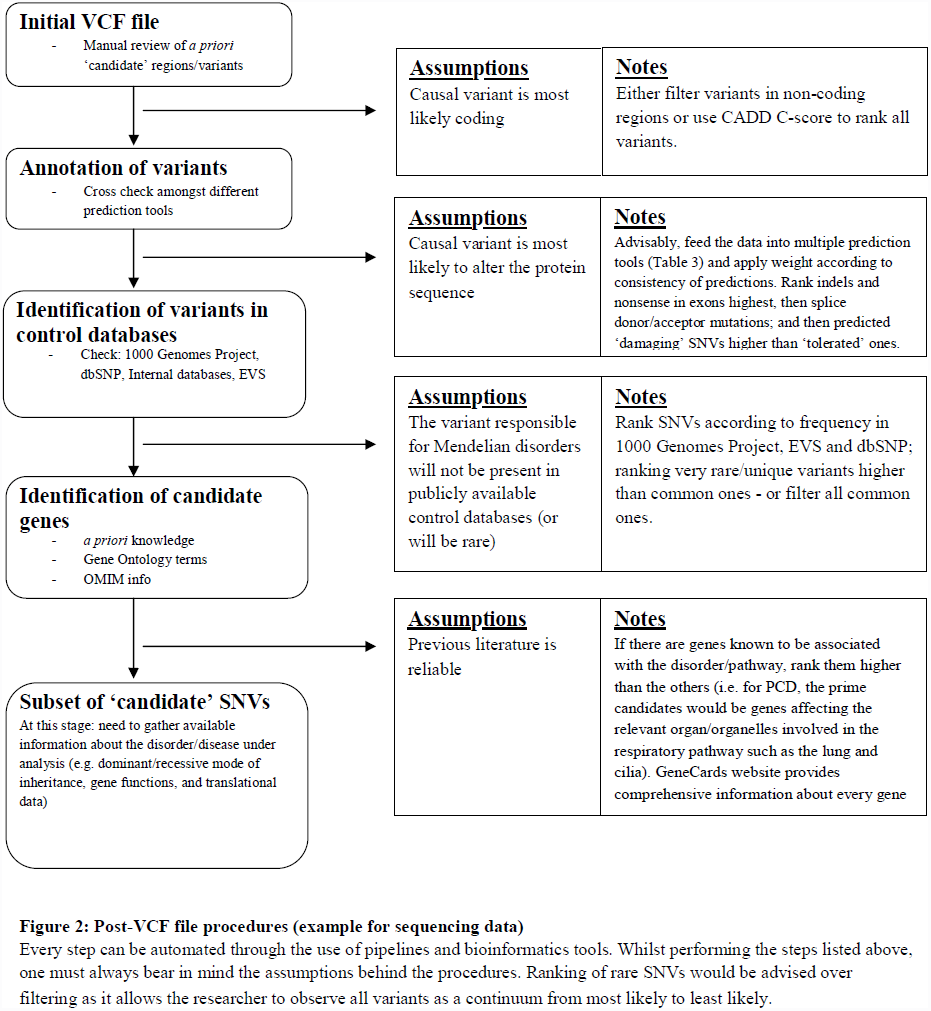
Post-VCF file procedures.

For individuals of European ancestry, a VCF file will have between eighty and ninety thousand variants for WES (more for individuals with African ancestry [35]); and approx. a tenth will be variants with ‘predicted high impact’ (also known as Φ variants i.e. rare nonsense, missense, splice-site acceptor or donor variants, exonic indels [36]). There are many algorithms which predict the functional effect of these variants (Table 3). A large proportion of these algorithms utilize sequence conservation within a multiple sequence alignment (MSA) of homologous sequences to identify intolerant substitutions, e.g. a substitution falling within a conserved region of the alignment is less likely to be tolerated than a substitution falling within a diverse region of the alignment (see Ng for a review [37]). A handful of these algorithms also utilize structural properties, such as the protein secondary structure and solvent accessible surface area, in order to boost performance. Well known examples of a sequence-based and structure-based algorithm are SIFT [38] and PolyPhen [39] respectively. Newer software such as FATHMM [40] and MutPred [41], which use state-of-the-art hidden Markov models and machine learning paradigms, are worth using for their performance. There are also several tools such as CONDEL-2 [42] which combine the output of several prediction tools to produce a consensus deleteriousness score. Although SIFT and Polyphen are highly cited tools, comparative analyses carried out by Thusberg *et al* and Shihab *et al* found FATHMM, MutPred and SNPs&GO to perform better using the VariBench benchmarking dataset containing missense mutations [40,43]. For predicting the effects of non-coding variants, GWAVA [12] and CADD [11] should be used. Also Human Splice Finder (latest: v3.0) can be used for intronic variants which predicts whether splicing is affected by the variant or not [44]. Many of these tools can be incorporated into the analyses through the Ensembl website (http://www.ensembl.org/info/docs/tools/vep/index.html) where VCF files are annotated [45].

**Table 3:** Tools for predicting variant effects.

Identifying neutral and pathogenic mutations
| Name | Reference | MCC | Comments |
| --- | --- | --- | --- |
| *SIFT | [84,85] | 0.30<br>(unweighted) | Highly cited with many projects using and citing it since 2001. Uses available evolutionary information and is continually updated. Easy to use through VEP. Provides two classifications: 'Deleterious' and 'Tolerated'. |
| *PolyPhen-2 | [39] | 0.43 | Provides a high quality multiple sequence alignment pipeline and is optimized for high-throughput analysis of NGS data. Cited and used by many projects of different types. Easy to use through VEP. Provides three classifications: 'Probably Damaging', 'Possibly Damaging' and 'Benign'. |
| *FATHMM | [40] | 0.72 | A highly performing prediction tool. Clear examples are available on the website. Offers flexibility to the user for weighted (trained using inherited disease causing mutations) and unweighted (conservation-based approach) predictions. Also offers protein domain-phenotype association information. |
| GERP++<br>(and GERP) | [86-88] | N/A | Determines constrained elements within the human genome; therefore variants in them are likely to induce functional changes. Can provide unique details about the candidate variant(s). |
| PhyloP | [89] | N/A | Helps detect non-neutral substitutions. Similar aim with GERP |
| CADD | [11] | - | Provides annotation and scores for all variants in the genome considering a wide range of biological features |
| GWAVA | [12] | - | Provides predictions for the non-coding part of the genome. |
| *SNAP | [90] | 0.47 | Predicts the effects of non-synonymous polymorphisms. Cited and used many times; and should be used to check whether the predicted effect is matched by the putative causal variant. However it was labelled 'too slow' for high throughput analyses by [43]. |
| PupaSuite | [91] | - | Identifies functional SNPs using the SNPeffect [92] database and evolutionary information. |
| Mutation Assessor-2 | [93] | - | Predicts the impact of protein mutations. User friendly website and accepts many formats. |
| *PANTHER | [94,95] | 0.53<br>(unweighted) | Predicts the effect of amino acid change based on protein evolutionary relationships. It provides a number ranging from 0 (neutral) to -10 (most likely deleterious) and allows the user to decide on the "deleteriousness" threshold. It is constantly updated making it a very reliable tool. |
| CONDEL-2 | [42] | - | Combines FATHMM and Mutation Assessor (as of version 2) in order to improve prediction. It theoretically outperforms the tools it is using in comparison to when the tools are used individually. |
| *MutPred | [41] | 0.63 | Predicts whether a missense mutation is going to be harmful or not based on a variety of features such as sequence conservation, protein structure and functional annotations. Praised in recent comparative study by [43]. |
| *SNPs&GO | [96] | 0.65 | Reported to have performed best amongst many prediction tools in [43]. Provides two classifications: 'Disease related' and 'neutral'. |
| Human Splicing Finder | [44] | N/A | Predicts the effect of non-coding variants in terms of alteration of splicing. Useful for compound heterozygotes if one allele is intronic. |
| Others | [97], [98], [99], [100] | 0.19<br>0.43<br>0.40<br>- | *nsSNPAnalyzer (requires 3D structure coordinates), *PhD SNP, *Polyphen (not supported any more), PMUT |

Many methods have been developed to predict the effect of missense mutations. Many of the tools listed above use different features and datasets to predict these effects; thus once the decision is made about which tool to use, the theory behind the predictions should always be kept in mind. Tools such as CONDEL-2 combine several of these tools to determine a consensus score which theoretically results in higher accuracy when compared to the individual tools.
\*Comprehensive information about the prediction tool including accuracy, specificity and sensitivity available in [43] and [40]. N/A: not applicable. MCC: Matthew's Correlation Coefficient. MCCs from [40].

### Further filtering/ranking

With current knowledge, there are fifty synonymous mutations with proven causality – complex traits and Mendelian disorders combined [46]. This is a very small proportion when compared to the thousands of published clinically relevant non-synonymous (i.e. missense and nonsense) mutations. Therefore, when filtering variants for rare monogenic disorders, not taking non coding variants and synonymous variants into account in the initial stages is a pragmatic choice. If ranking is preferred, then tools such as SilVA [47] which ranks all synonymous variants and CADD [11] which ranks all variants (including synonymous variants) in the VCF files should be used.

Highly penetrant (Mendelian or common-complex) disease causal variants are expected to be very rare, therefore most of them should not appear in publicly available datasets. However filtering all variants present in dbSNP which is common practice, should not be carried out as amplification and/or sequencing errors as well as potentially causal variants are known to make their way into this database [48, 49]. Thus use of a MAF threshold (e.g. ≤ 0.1% in 1000 Genomes and/or EVS) is a wiser choice in contrast to using absence in dbSNP as a filter. Upon completion of these steps, a smaller subset of variants with strong candidacy will remain for further follow up to determine causality.

As many online tools are expected to keep logs of the processes undergoing in their servers, to protect confidentiality of genetic information downloading a local version of the chosen tools (or the VEP cache from the Ensembl website) is recommended. VEP also enables incorporation of MAF from the EVS and the 1000 Genomes Project – and many other annotations (e.g. conservation scores, is variant position present in HGMD public version, PubMed), which will make the filtering steps more manageable.

## STAGE 3 – BUILDING EVIDENCE FOR CAUSALITY

Figure 3 suggests the route to take to help differentiate causal variant(s) from non-causal ones for Mendelian disorders. At this stage one must gather all information that is available about the disorder and use them to determine which inheritance pattern fits the data and what complications there might be (e.g. the possibility of compound heterozygotes in disorders which show allelic heterogeneity). Supp. Figure S2 can be used to observe the contrast between the routes taken when analysing Mendelian (Figure 3) and complex disorders.

**Figure 3:**
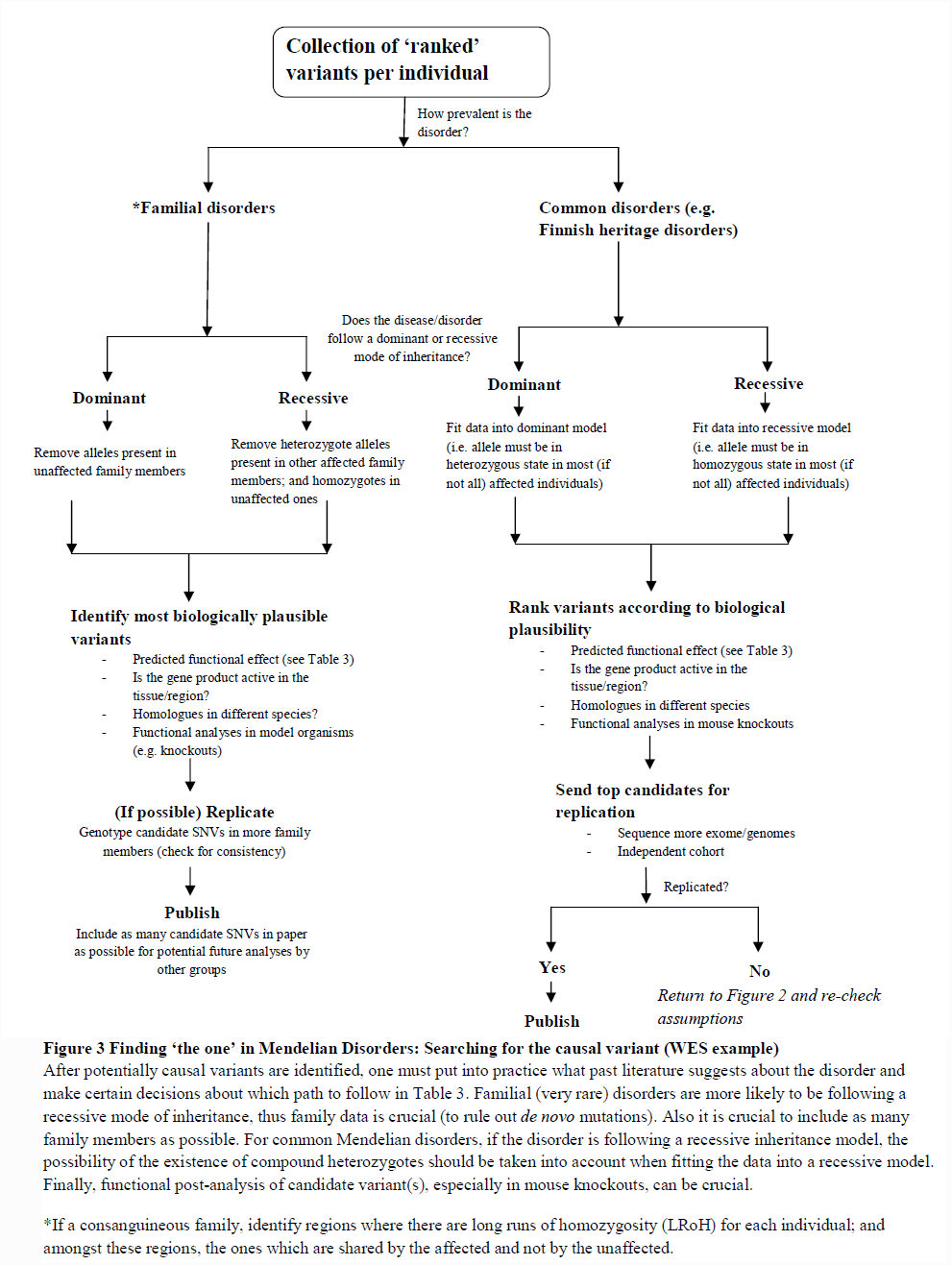
Finding ‘the one’ in Mendelian Disorders.

### Public data as a source of evidence

Having a candidate gene list based on previously published literature (e.g. by using OMIM or disease/pathway specific databases such as the Ciliome database [50]) and knowledge about the biology of the disorder (e.g. biological pathways) is useful. Software such as STRING and KEGG predicts protein-protein interactions using a variety of sources [51, 52]. SNPs3D is a user friendly interface which is designed to suggest candidates for different disorders [53]. UCSC Gene Sorter (accessible from https://genome.ucsc.edu/) is another useful tool for collating a candidate gene list as it groups gene according to several features such as protein homology, coexpression and gene ontology (GO) similarity. Uniprot’s (http://www.uniprot.org/) Blast and Align functions can provide essential information about the crucial role a certain residue plays within a protein if it is highly conserved throughout many species. This is especially important for SNVs (excluding nonsense mutations as they truncate the protein) where the SNV itself should be causal.

**Figure 4.**
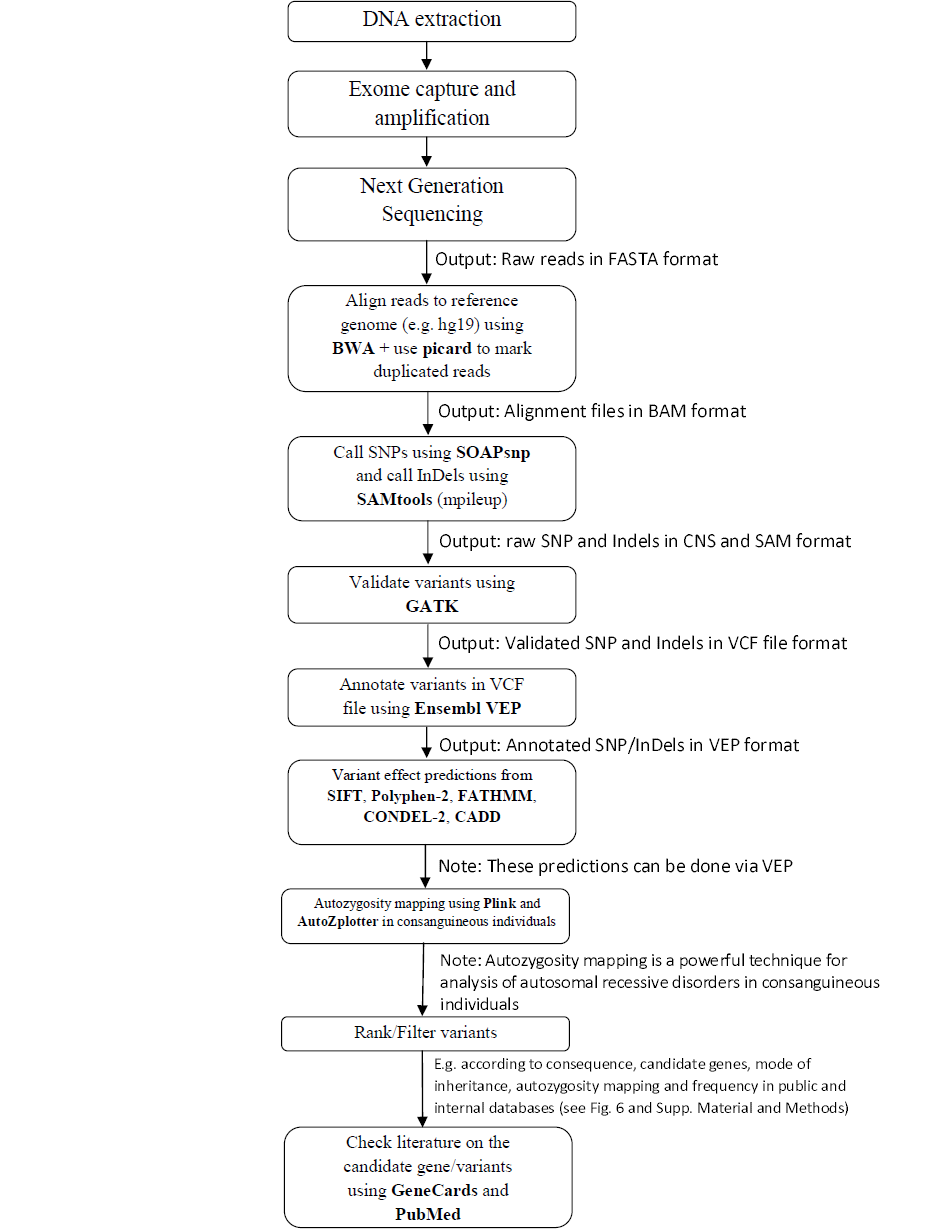
Summary of whole analysis process: DNA sample to identification of variant. The tools mentioned here are the ones we prefer to use for a variety of reasons such as documentation, ease of use, performance, multi-platform compatibility and speed. See Supp. Material and Methods for examples of parameters/commands to use where applicable.

**Figure 5:**
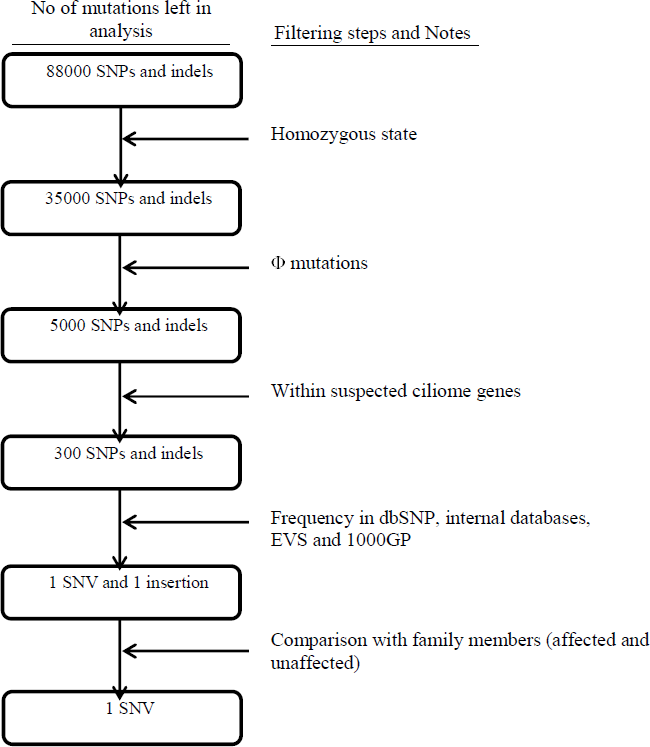
Filtering steps applied to all mutations in the exome (Primary ciliary dyskinesia example) After all the filtering steps in the above figure are applied, the total will be reduced to a single candidate. The numbers here are for illustration purposes only (adapted from [36]). Homozygosity step is added as PCD is an autosomal recessive disorder. Φ mutations are ‘predicted high impact’ mutations as proposed by Alsaadi and Erzurumluoglu *et al* [36] (see SO_terms_SNP.txt in Supp. data).

An example of the filtering process for an autosomal recessive disorder such as PCD is depicted in Figure 5. If several variants pass the filtering steps, information about the relevant genes should be gathered using databases such as GeneCards (www.genecards.org/) and NCBI Gene (www.ncbi.nlm.nih.gov/gene) for functional information, GEO Profiles (www.ncbi.nlm.nih.gov/geoprofiles) and Unigene (www.ncbi.nlm.nih.gov/unigene) for translational data about the gene’s product; and if available, one can check if a homologue is present in different species using databases such as HomoloGene (www.ncbi.nlm.nih.gov/homologene) and whether a similar phenotype is observed in model organisms. For example, if the disorder affects the cerebral cortex but the gene product is only active in the tissues located in the foot, then one cannot make a good argument about the identified variant in the respective gene as being ‘causal’. There are many complications that may arise depending on the disorder such as genetic (locus) heterogeneity [54], allelic heterogeneity [55] and incomplete penetrance [56]. Therefore gathering as many cases from the same family is helpful. However for very rare Mendelian disorders this may not be possible, thus it is important to seek other lines of evidence (e.g. animal models, molecular analyses).

### Mapping causal loci within families

For rare Mendelian disorders, familial information can be crucial. The availability of an extended pedigree can be very informative in mapping which variant(s) fits the mode of inheritance in the case(s) and not in the unaffected members of the family (e.g. for autosomal recessive mutations, confirming heterozygosity in the parents is a must). This will provide linkage data where its importance is best displayed by Sobreira *et al* where WES data from a single proband was sufficient in discovering the causal variants in two different families [57]. Where available, previously published linkage data (i.e. associating a chromosomal region to a Mendelian disorder) should also made use of.

Traditionally a LOD score of 3 (Prob. = 1/1000) is required for a variant/region to be accepted as causal. Reaching this threshold requires many large families with many affected individuals. However this is not feasible for most disease causal variants (which are very rare by nature) and other lines of evidence such as animal knockouts, molecular studies and alignments are required to make a case for the causality of variants, especially mutations which are not stop gains (e.g. missense).

As mentioned previously, understanding the characteristics of a Mendelian disorder is important. If the disorder is categorised as ‘familial’ (i.e. occurs more in families than by chance alone), which are usually very rare by nature, then availability of familial data becomes crucial – as unaffected members of the family are going to be the main source of information when determining neutral alleles. Any homozygous (and rare) stop gains in previously identified genes would be prime candidates.

Approach taken in families is different from the approaches taken when analysing common Mendelian disorders using unrelated individuals. For common Mendelian disorders (e.g. Finnish Heritage disorders [58-60]), fitting the dataset into a recessive inheritance model requires most (if not all) affected individuals to have two copies of the disease allele, enabling the identification of founder mutations as they will be overrepresented in the cases. These variants will be homozygous through endogamy and not consanguinity.

#### Autozygosity mapping

For consanguineous subjects, the causal mutation usually lies within an autozygous region (characterised by long regions of homozygosity, LRoH, which are generally >5Mb, see [61]), thus checking whether any candidate genes overlaps with an LRoH can narrow region(s) of interest. There are several tools which can identify LRoHs such as Plink, AutoSNPa and AgilentVariantMapper. We have made available a python script (AutoZplotter) to plot heterozygosity/homozygosity status of variants in VCF files to allow for screening of short autozygous regions as well as LRoHs.

#### AutoZplotter

There are several software which can detect long runs of homozygosity reliably (>5Mb), however they struggle to identify regions that are shorter than these. Therefore we developed AutoZplotter which plots homozygosity/heterozygosity state and enables quick visualisation of suspected autozygous regions. The input format of AutoZplotter is VCF thus it suits any type of genetic data (e.g. SNP array, WES, WGS). AutoZplotter was used for this purpose in a previous study by Alsaadi *et al* [18].

### Exceptional cases

There can always be exceptional cases (in consanguineous families also) such as compound heterozygotes (i.e. individuals carrying different variants in the two copies of the same gene). This would require haplotype phasing and the confirmation of variant status (i.e. heterozygosity for one allele and absence of the other) in the parents and the proband(s) by sequencing of PCR amplicons containing variant or genotyping the variant directly. Beagle and HAPI-UR are two widely used haplotype phasing tools for their efficiency and speed [62, 63].

### Identifying highly penetrant variants for common-complex disorders

For common complex disorders, identifying causal variants in outbred populations has proven to be a difficult and costly process (Supp. Figure S2); and these disorders can have many unknowns such as the significance of environmental factors [64-66] and epistasis [67]. Many of the causal variants may be relatively rare (and almost always in heterozygous state) in the population introducing issues with statistical power. Traditional GWAS do not attempt to analyse them thus they are largely ignored – leaving a lot of heritability of common complex disorders unexplained. Analysing individuals with extreme phenotypes where the segregation of disease mimics autosomal recessive disorders (e.g. in consanguineous families) can be useful in identifying highly penetrant causal genes/mutations for complex disorders (e.g. obesity and leptin gene mutations [68]). The genetic influence in these individuals is predicted to be higher and are expected to have a single highly penetrant variant in homozygous state. These highly penetrant mutations can mimic Mendelian disorders causal variants. Therefore similar study designs can be used (e.g. Autozygosity/homozygosity mapping).

## CONCLUSIONS

The NGS era has brought data management problems to traditional geneticists. Many data formats and bioinformatics tools have been developed to tackle this problem. One can easily be lost in the plethora of databases, data formats and tools. “Which tools are out there? How do I use it? What do I do next with the data I have?” are continually asked questions. This review aims to guide the reader in the rapidly changing and ever expanding world of bioinformatics. Figure 4 depicts a summary of the analysis process from DNA extraction to finding the causal variant, putting into perspective which file formats are expected at each step and which bioinformatics tools we prefer due to reasons mentioned before. Researchers can then appreciate the stage that they are at and how many other steps are required for completion as well as knowing what to do at each step.

Whole exome sequencing is the current gold standard in the discovery of highly penetrant disease causal mutations. As knowledge on the non-coding parts of the genome can still be considered to be in its early days, the human exome is still a pragmatic target for many. As approx. 1600 known Mendelian disorders (and ∼3500 when suspected ones are included) and most common-complex disorders are still waiting for their molecular basis to be figured out (from omim.org/statistics/entry, true as of 15/07/14), future genetic studies have much to discover. However for these projects to be fruitful, careful planning is needed to make full use of available tools and databases (see Table 4).

**Table 4:** What is needed for a genetic study?

| **Material** | **Notes** |
| --- | --- |
| 'Sufficient' number of high-quality sequencing/genotype data | Amount needed can vary from one proband and a few family members (for very rare Mendelian disorders) to 10000 case and controls (for certain complex disorder/traits) |
| List of candidate genes | Websites such as <u>OMIM</u> and <u>GHR</u>; and software such as SNPs3D can be helpful |
| Identification of variant calling tool | Such as in Table 2 |
| Identification of variant effect predictor tool | Such as in Table 3; tools usually require conversion of VCF to VEP format (Ensembl website) |
| Knowledge of human population variation databases | i.e. HapMap, 1000 Genomes Project, EVS, dbSNP, internal databases |
| Knowledge of databases storing information about genes and their products | i.e. OMIM, Gene (NCBI), GeneCards, Unigene (NCBI), GEO Profiles (NCBI), HomoloGene (NCBI), Mouse knockout databases (such as <u>MGI</u>, <u>TIGM</u> and <u>NC3RS</u>). Search the literature using PubMed and/or Web of Science. |

The most important factors when carrying out a genetic association study are (i) the availability of data (ii) expertise and (iii) careful planning

Finally, with this paper we have also made AutoZplotter available (input format: VCF), which plots homozygosity/heterozygosity state and enables quick visualisation of suspected autozygous regions. This can be important for shorter autozygous regions where other autozygosity mappers struggle.

## Acknowledgements

I thank Tom G. Richardson, Dr. Hashem Shihab, Denis Baird, Dr. Tom Gaunt, Prof. Ian Day and Dr. Santi Rodriguez for their advice and encouragement.

### Funding

Mesut Erzurumluoglu is a PhD student funded by the Medical Research Council (MRC UK).

## Supp. Material and Methods

These commands are here to guide the user. However where complications arise, other options may have to be included thus requires reading documentation provided by the bioinformatics tools.

**Parameters used in BWA for read alignments:** bwa aln -o 1 -e 50 -m 10000 -t 4 -i 15 -q 10 -I

*-I at the end is for Illumina NGS platforms*

**Parameters used in GATK (for SNPs):** java -jar GenomeAnalysisTK.jar -T UnifiedGenotyper - stand_call_conf 50 -stand_emit_conf 10.0 -A DepthOfCoverage -A RMSMappingQuality -baq CALCULATE_AS_NECESSARY

**Parameters used in GATK (for InDels):** java -jar GenomeAnalysisTK.jar -T UnifiedGenotyper - stand_call_conf 50 -stand_emit_conf 10.0 -A DepthOfCoverage -A RMSMappingQuality -baq CALCULATE_AS_NECESSARY -glm INDEL

**Obtaining Ensembl VEP annotations for VCFs (including SIFT, Polyphen and Condel predictions):**

1. Download latest package (and *plugins) from Ensembl website: (www.ensembl.org/info/docs/variation/vep/index.html)
2. Tar xvf downloaded file(s)
3. perl INSTALL.pl – and download *Homo sapiens* cache(s)
4. perl variant_effect_predictor.pl -i **file.vcf** -o **file.vep** --protein --cache --regulatory --gmaf --force_overwrite --sift b --polyphen b -- plugin Condel,/data/home/∼/ensembl-tools-release-75/scripts/variant_effect_predictor/ensembl-variation-VEP_plugins-e6cec6a/config/Condel/config,b --fork 8 --canonical --individual all --pubmed --maf_esp --symbol

*to use Condel plugin:

1- Download latest Ensembl plugins from: https://github.com/ensembl-variation/VEP_plugins

2- tar -xvf downloaded file

2- mv Condel.pm ∼/.vep/Plugins (create Plugins folder if not there; also .vep is a hidden folder)

3- edit the condel_SP.conf file (in config/Condel/config/) and set the ‘condel.dir’ parameter to /data/home/∼/variant_effect_predictor/ensembl-variation-VEP_plugins-e6cec6a/config/Condel

**Example of commands used to filter variants in VEP file:** To grab list of all rare/unique and homozygous mutations in candidate genes: grep -f **Candidate_genes.txt file.vep** | grep -f **SO_terms_SNP.txt** | grep CANONICAL | grep HOM | grep _[A-Z]/ > **file_candidate_mutations.txt**

or use grep GMAF=[A-Z]:0.00 instead of grep _[A-Z]/ for variants which are present in the 1000GP but rare

**Files used:**

**Candidate_genes.txt:** a text file containing Ensembl IDs of your candidate genes – one per row

**SO_terms_SNP.txt:** a text file containing VEP SO terms which would be classified as a Φ mutation (available as Supp. File)

**Command for Autozygosity plotting in AutoZplotter:** python autozplotter.py

**Parameters used for Autozygosity mapping in Plink:** plink --file <ped/map> --homozyg --noweb -- homozyg-window-kb 1000 --homozyg-window-het 1 --homozyg-group --out <output>

**Supp. Figure S1.**
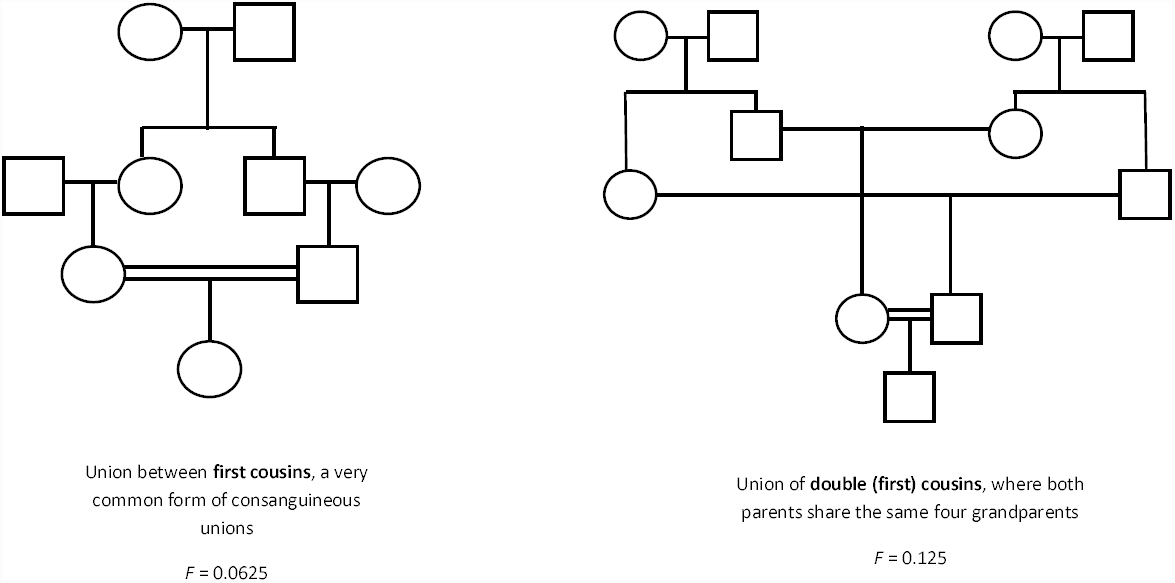
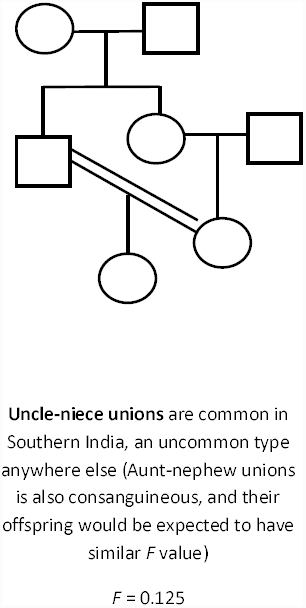
Consanguineous Unions where F ≥ 0.0625

**Supp. Figure S2.**
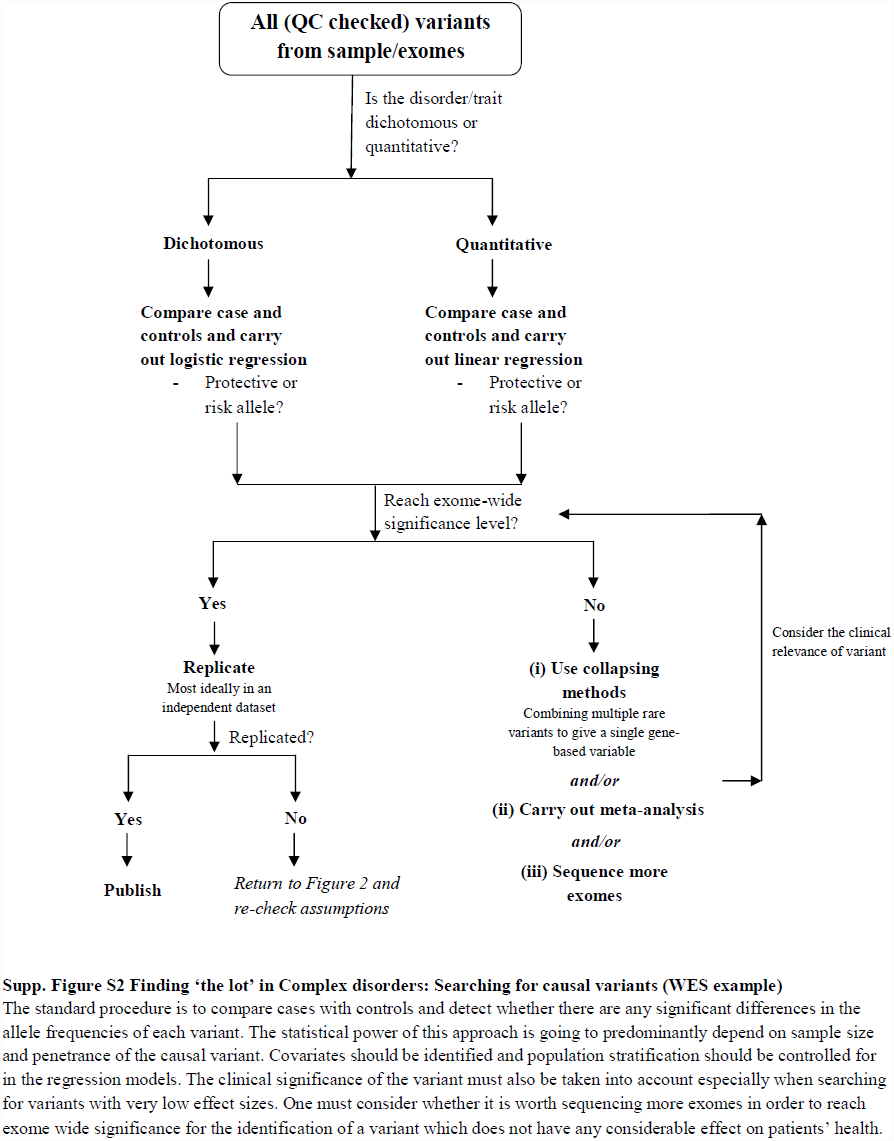
Finding ‘the lot’ in Complex disorders

